# Modulation of Actin network and Tau phosphorylation by HDAC6 ZnF UBP domain

**DOI:** 10.1101/702571

**Authors:** Abhishek Ankur Balmik, Shweta Kishor Sonawane, Subashchandrabose Chinnathambi

**Author notes:** The authors wish it to be known that, in their opinion, the first two authors should be regarded as joint First Authors. To whom correspondence should be addressed: **Prof. Subashchandrabose Chinnathambi**, Neurobiology group, Division of Biochemical Sciences, CSIR-National Chemical Laboratory (CSIR-NCL), Dr. Homi Bhabha Road, 411008 Pune, India. **Email:**.

## Abstract

Microtubule-associated protein Tau undergoes aggregation in Alzheimer’s disease and a group of other related diseases collectively known as Tauopathies. In AD, Tau forms aggregates, which are deposited intracellularly as neurofibrillary tangles. HDAC6 plays an important role in aggresome formation where it recruits polyubiquitinated aggregates to the motor protein dynein. Here, we have studied the effect of HDAC6 ZnF UBP on Tau phosphorylation, ApoE localization, GSK-3β regulation and cytoskeletal organization in neuronal cells by immunocytochemistry. Immunocytochemistry reveals that HDAC6 ZnF UBP can modulate Tau phosphorylation and actin cytoskeleton organization when the cells are exposed to the domain. HDAC6 ZnF UBP treatment to cells does not affect their viability and resulted in enhanced neurite extension and formation of structures similar to podosomes, lamellipodia and podonuts suggesting its role in actin re-organization. Also, HDAC6 treatment showed increased nuclear localization of ApoE and tubulin localization in microtubule organizing centre. Our studies suggest the regulatory role of this domain in different aspects of neurodegenerative diseases.

## Introduction

Tau pathology is implicated in several neurodegenerative diseases including Alzheimer’s and Parkinson’s disease, Progressive supranuclear palsy, Pick’s disease, Corticobasal degeneration and Post-encephalitic parkinsonism [1,2]. Tau is comprised of two N-terminal inserts, a poly-proline domain and four imperfect repeat domains (Fig. 1A). The four repeat regions are the functional and pathological unit of Tau as it is involved in microtubule-binding as well as aggregate formation. In Alzheimer’s disease and related Tauopathies, Tau gets detached from microtubules and associates with other Tau molecules to form intracellular aggregates in the form of NFTs [3,4]. Tau acquires pathological form either through the genetic predisposition on chromosome 17 termed as FTDP-17 or hyperphosphorylation and several other post-translational modifications [5,6]. Phosphorylation is a well-studied PTM of Tau responsible for its conversion to pathological form in AD condition [7–9]. GSK-3β is one of the major kinase involved in Tauopathies [10]. Tau is phosphorylated by GSK-3β on both primed (after pre-phosphorylation) and unprimed sites and affects its ability to bind and stabilize microtubules [11,12]. In Alzheimer’s disease, the structure and function of Tau protein is modified by various cellular and molecular factors such as oxidative stress, post-translational modifications and interaction with its binding partners [13]. This result in the accumulation of Tau protein aggregates as intracellular inclusion bodies called neurofibrillary tangles (NFTs). The abnormal accumulation of Tau as NFTs affects various cellular processes including microtubule dynamics and intracellular transport mechanism dependent on the cytoskeletal networks.

**Figure 1.**
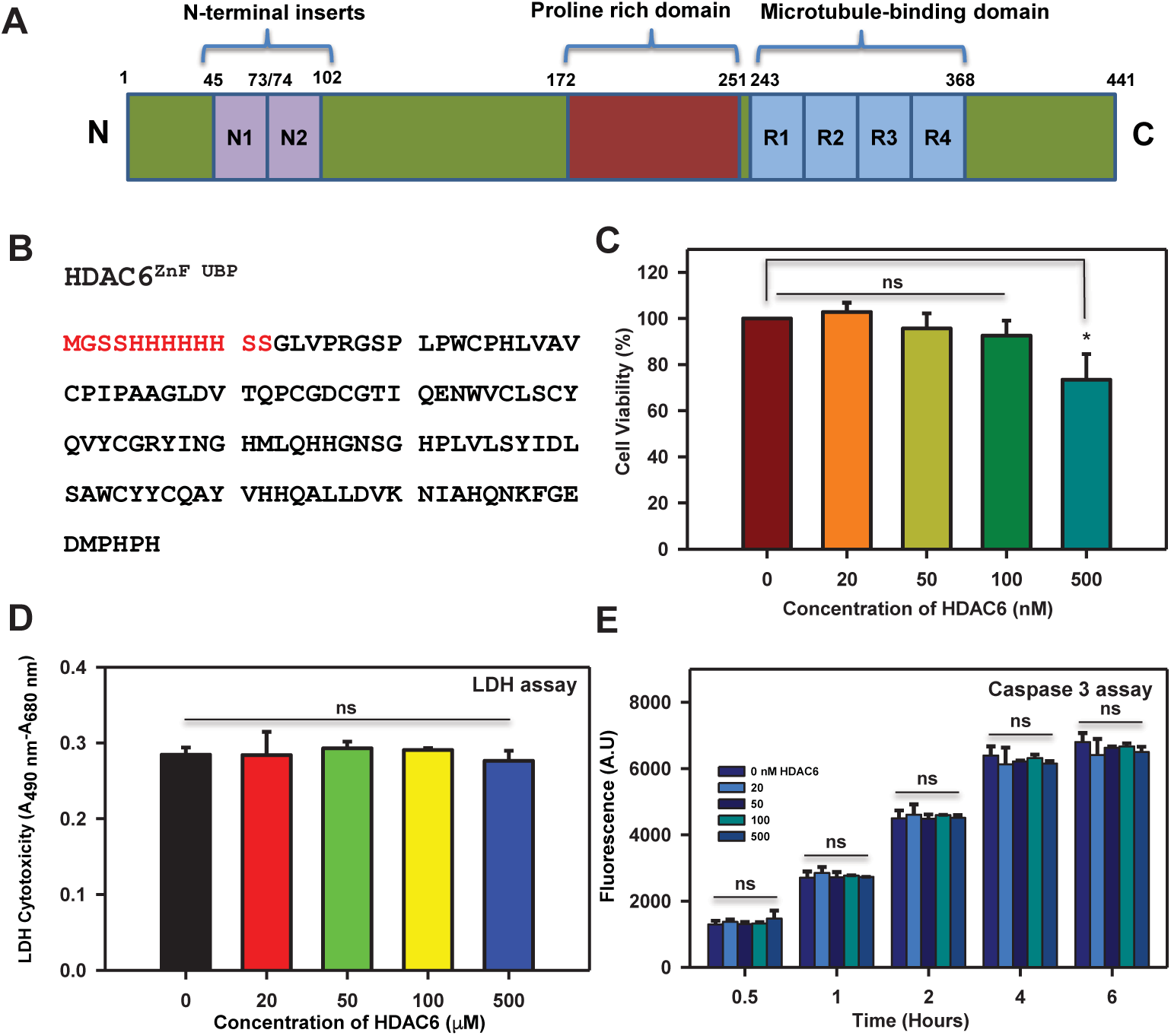
HDAC6 ZnF UBP treatment to neuro2a cells does not have any toxic effect. A) Bar diagram for the human microtubule-associated protein Tau comprising of two N-terminals inserts and four repeat regions preceded by a polyproline domain. B) Amino acid sequence of human HDAC6 Zinc finger ubiquitin-binding domain. This domain is located on the C-terminal of HDAC6 and associates with polyubiquitinated protein aggregates to mediate the formation of aggresomes. C) MTT assay was carried out to determine the viability of neuro2a upon HDAC6 ZnF UBP treatment. Neuroblastoma cells treated with HDAC6 at different concentrations show minimum toxicity and maintain viability at 80% at highest concentration of 500 nM. D) LDH release assay determines the damage to the cell membrane upon exposure to test molecule. The membrane leakage assay (LDH assay) shows that HDAC6 does not disrupt the cell membrane and affect cell viability. E) Apoptosis rate of neuro2a cells upon HDAC6 ZnF UBP treatment was assayed using Caspase-3 assay. Caspase-3 assay shows increased levels of caspase-3 with successive time interval but do not differ from control samples. (ns – non-significant, * indicates P ≤ 0.05).

The proteins upon misfolding are targeted for degradation by ubiquitination and these poly-ubiquitinated protein aggregates are degraded and recycled in proteasome [14–17]. Aggregation and subsequent accumulation of Tau protein aggregates are primarily tackled by chaperones and ubiquitin proteasomal system. In case of disruption of ubiquitin proteasome system (UPS), the aggregates are directed towards the formation of aggresomes which serves as the cytoprotective response upon the failure of UPS [17–20]. HDAC6 is a class II histone deacetylase mainly present in the cytoplasm involved in the regulation of various cellular functions. It consists of two catalytic deacetylase domains and a unique ZnF UBP (Zinc finger ubiquitin binding protein) domain, which sets it apart from other HDACs [21–23]. The proteins upon misfolding are targeted for degradation by ubiquitination and these poly-ubiquitinated protein aggregates are degraded and recycled in proteasome [14–17]. Aggregation and subsequent accumulation of Tau protein aggregates are primarily tackled by chaperones and ubiquitin proteasomal system. In case of disruption of ubiquitin proteasome system (UPS), the aggregates are directed towards the formation of aggresomes which serves as the cytoprotective response upon the failure of UPS [17–20]. One of the major functions of HDAC6 is recruiting the polyubiquitinated protein aggregates to Dynein/Dynactin complex and carrying them to MTOC for aggresome formation. HDAC6 acts on cortactin and mediates its association with F-actin to facilitate cell motility [24]. The function of HDAC6 in both UPS and autophagy indicate its role as a possible link between the two mechanisms [25–27]. The impairment of UPS function acts as a cue for the activation of a compensatory mechanism for the clearance of protein aggregates. It has been studied in the *Drosophilla* model of spinobulbar muscular atrophy, where expression of HDAC6 has been found to effectively cause rescue from UPS impairment induced neurodegeneration by triggering the autophagic clearance of protein aggregates [26]. In another study, HeLa cells transfected with PolyQ Huntingtin forms intracellular protein aggregates, which require HDAC6 for autophagic clearance after inhibition of proteasomal system [28]. Overall, the function of HDAC6 with respect to protein aggregate clearance and autophagy induction serves as a protective mechanism. The HDAC6 expression level increases sharply in protein misfolding diseases [29].

Tau exists in hyperphosphorylated state in NFTs. Alzheimer’s disease is associated with the upregulation of cellular kinases like GSK-3β and CDK5 [30]. There a number of serine and threonine residues in repeat region of Tau which are phosphorylated in AD through proline directed kinases [31]. On the other hand protein phosphatases like PP1 and PP2A are known to be downregulated in AD failing to reverse the effect of hyperphosphorylation [32–34]. Thus, there is an imbalance between the kinase and phosphatase function in neurons. Another important aspect of neurodegenerative diseases involve the mis-functioning of its cytoskeletal elements. Microtubule network and actin organization were found to be distorted leading to impaired cellular trafficking and other associated functions [35,36]. Actin organization is crucial for synaptic signalling in neurons where they are involved in the formation of dendritic spines for neurotransmission. Impaired actin assembly and depolymerisation leads to loss of dendritic spines ultimately causing neuronal death [37,38].HDAC6 is a key protein in the regulation of both actin and microtubule organization through its deacetylase activity [39]. However, the role of its ZnF UBP domain in cytoskeletal function is lesser studied.In the present study, we studied the role of HDAC6 ZnF UBP domain (Fig.1B) in modulating different cellular events such as phosphorylation, cytoskeletal assembly and cellular localization. HDAC6 ZnF UBP domain was found to affect actin and tubulin organization, Tau phosphorylation and localization of ApoE in neuronal cells. Enhancement in podosome and lamellipodia-like structures were found when neuronal cells were exposed to HDAC6 ZnF UBP domain suggesting its direct role in actin organization. HDAC6 ZnF UBP treatment also resulted in enhanced tubulin localization in MTOC indicating its possible role in tubulin polymerization events. Our findings suggest the role of HDAC6 ZnF UBP as the direct modulator of Tau phosphorylation and cytoskeletal dynamics.

## RESULTS

### HDAC6 ZnF UBP is non-toxic and non-apoptotic

Neuronal cells were exposed to HDAC6 ZnF UBP exogenously to observe the effect on different cellular functions. In order to study the effects of HDAC6 ZnF UBP, a minimum toxic dose was determined by viability assay as well as membrane leakage assays. Neuronal cells were treated with a range of concentrations of HDAC6 ZnF UBP (20-500 nM). The HDAC6 internalization was monitored for 20-500 nM concentrations by immunostaining for HDAC6 and anti-His-tag (Supp. Fig. 1). The cell viability carried out by MTT assay showed no decrease in viability. The cell viability was maintained at 80% even in highest HDAC6 ZnF UBP concentration of 500 nM (Fig.1C). Similarly, LDH assay was carried out to check the effect of HDAC6 on membrane integrity in terms of LDH release. Neuronal cells showed intact membrane integrity when treated with HDAC6 ZnF UBP in 20-500 nM concentration range (Fig.1D). Thus, HDAC6 ZnF UBP did not show toxicity in neuroblastoma cells. Further to confirm the non-toxic nature of HDAC6 ZnF UBP in neuronal cells we studied the apoptosis on treatment with HDAC6 ZnF UBP by caspase-3 assay. Cells undergo apoptosis under stressful conditions-mediated by endoproteases called caspases. Caspase-3 is an executioner caspase responsible for DNA fragmentation and degradation of cytoplasmic proteins. The caspase-3 assay showed basal level of activity in all the treated and control samples. No difference was observed in the HDAC6 ZnF UBP treated and untreated control samples in terms of cell viability, suggesting that HDAC6 ZnF UBP treatment do not induce apoptosis in the neuro2a cells (Fig.1E). The preliminary studies on cell viability and morphology inferred that HDAC6 ZnF UBP treatment do not show cytotoxic effect on neuro2a cells. Hence, the cells were treated with a moderate concentration of 50 nM HDAC6 ZnF UBP for subsequent experiments.

### HDAC6 ZnF UBP enhances levels of pGSK-3β

Modulation of GSK-3β by HDAC6 ZnF UBP may involve regulation of Akt *via* PP1. PP1 dephosphorylates Akt which is a negative regulator of GSK-3β in its phosphorylated state [40]. HDAC6 is known to associate with PP1 through its C-terminal region which corresponds to ZnF UBP domain [41]. HDAC6 ZnF UBP addition to cells increases PP1-HDAC6 association rendering Akt in its active phosphorylated state, which in turn phosphorylates Ser9 on GSK-3β downregulating its activity (Fig. 2A). The level of GSK-3β in its phosphorylated and non-phosphorylated form determines its activity as a kinase. GSK-3β is known to associate with HDAC6 to counteract its function to induce LPS-tolerance in astrocytes [42]. We mapped GSK-3β and pGSK-3β (pSer9) levels by immunofluorescence in neuronal cells in basal conditions and upon HDAC6 ZnF UBP treatment. There was no marked difference in level of GSK-3β in HDAC6 treated and untreated cells (Fig.2B). However, immunofluorescence analysis of HDAC6 ZnF UBP treated cells showed significant increase in the pGSK-3β levels (Fig.2C). GSK-3β function in cells is regulated mainly by inhibitory phosphorylation at ser9 or ser21. The kinase activity of GSK-3β is reduced with phosphorylation at ser9 as it affects the binding of primed substrates with GSK-3β. The increase in pGSK-3β in neuronal cells signifies the reduced GSK-3β activity on primed Tau substrate [43]. HDAC6 ZnF UBP exposed to the cells resulted in increased levels of pGSK-3β (Ser9), which suggests reduced Tau phosphorylation. The increased levels of pGSK-3β suggest that GSK-3β activity is down-regulated upon HDAC6 ZnF UBP treatment.

**Figure 2.**
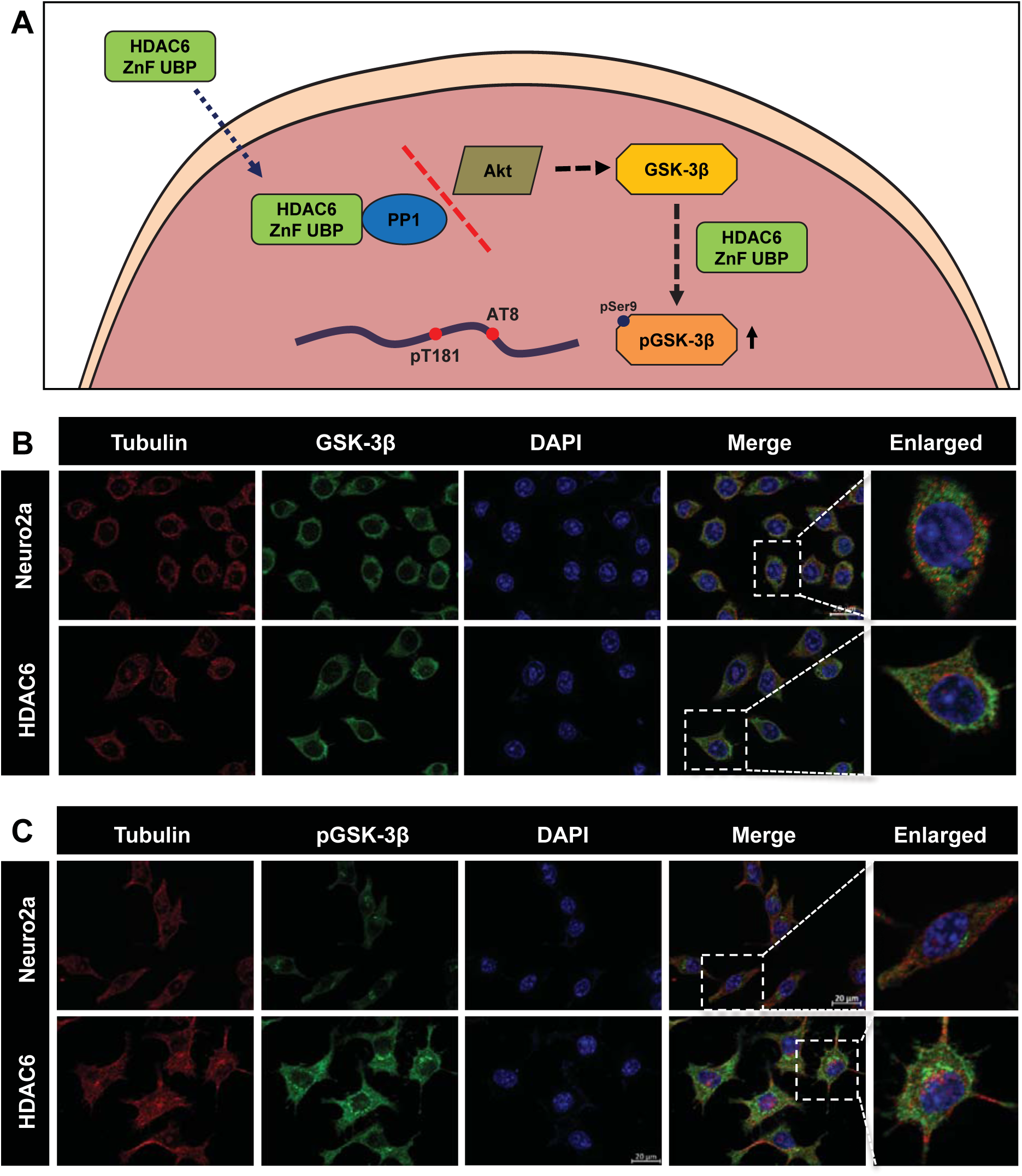
Downregulation of GSK-3β activity by HDAC6. The levels of GSK-3β and pGSK-3β reflect its activity in the cell. Enhanced levels of pGSK-3β were observed while total GSK-3β remained unaltered upon the treatment of HDAC6 ZnF UBP domain. A) Neuro2a mapped for total GSK-3β shows their unaltered levels upon HDAC6 ZnF UBP. B) The activity of GSK-3β is governed by phosphorylation at specific residues which either upregulates or downregulates its activity. Inhibitory phosphorylation of GSK-3β at Ser9 increases upon HDAC6 ZnF UBP treatment. The enlarged image shows the elevated level of pGSK-3β compared to neuro2a cell control. C) Proposed mechanism for the effect of HDAC6 ZnF UBP on the regulation of GSK-3β activity. GSK-3β activity is negatively regulated by Akt kinase, which phosphorylates GSK-3β at Ser9. Akt is active in its phosphorylated state. HDAC6 ZnF UBP associates with PP1 restricting it from dephosphorylating Akt, hence resulting in its active phosphorylated state.

### HDAC6 ZnF UBP reduces Tau phosphorylation in neuronal cells

Tau phosphorylation is the key event in the pathogenesis of Alzheimer’s disease. Tau phosphorylation is required for its function in microtubule interaction and stabilization. Under pathological conditions Tau becomes hyperphosphorylated due to imbalance in kinase and phosphatase level or activity [7-9,44] Okadaic acid (OA) was used as an inducer of hyperphosphorylation as it inhibits protein phosphatase 2A (PP2A), thus increasing the overall phosphorylation level [45]. Untreated control cells and HDAC6 ZnF UBP treated cells alone showed basal levels of phospho-Tau at epitopes pT181 and pS202/T205 (AT8). The levels of pT181 were increased in OA treated cells. Cells supplemented with HDAC6 ZnF UBP with OA showed lower levels of phospho Tau at T181 as compared to positive control (Fig.3A). Tau phosphorylated at pT181 was dominantly seen in the nucleus especially in the positive control (enlarged images). Similar results were observed for phospho Tau epitope AT8. The enlarged images clearly show the increased levels of phospho-AT8 in positive control as compared to OA+HDAC6 ZnF UBP treatment (Fig.3B). pT181 and AT8 are two of the crucial epitopes of pathological Tau in AD. HDAC6 ZnF UBP treatment lowered down the level of these two phospho-epitopes suggesting its possible role in modulating Tau phosphorylation. The level of both phospho-tau epitopes (pT181 and AT8) were quantified by their immunofluorescence levels in different experimental groups (No. of fields selected for quantification of pT181 and AT8, n = 5). Untreated cells (CC) showed no significant difference from HDAC6 treated or OA+HDAC6 treated in pT181 immunostaining. OA treatment resulted in increased pT181 levels, which was found to be reduced with HDAC6 ZnF UBP treatment (Fig. 3C). Similar results were obtained for quantification of AT8 immunostaining. HDAC6 ZnF UBP treatment along with OA reduced AT8 level, bringing it down similar to untreated cells (CC) when compared to OA alone treatment group. AT8 levels were also found to be lesser in HDAC6 ZnF UBP treatment as compared to untreated (Fig. 3D). Quantification data was analyzed by one way ANOVA followed by Tukey’s HSD test (significant at α = 0.05).

**Figure 3.**
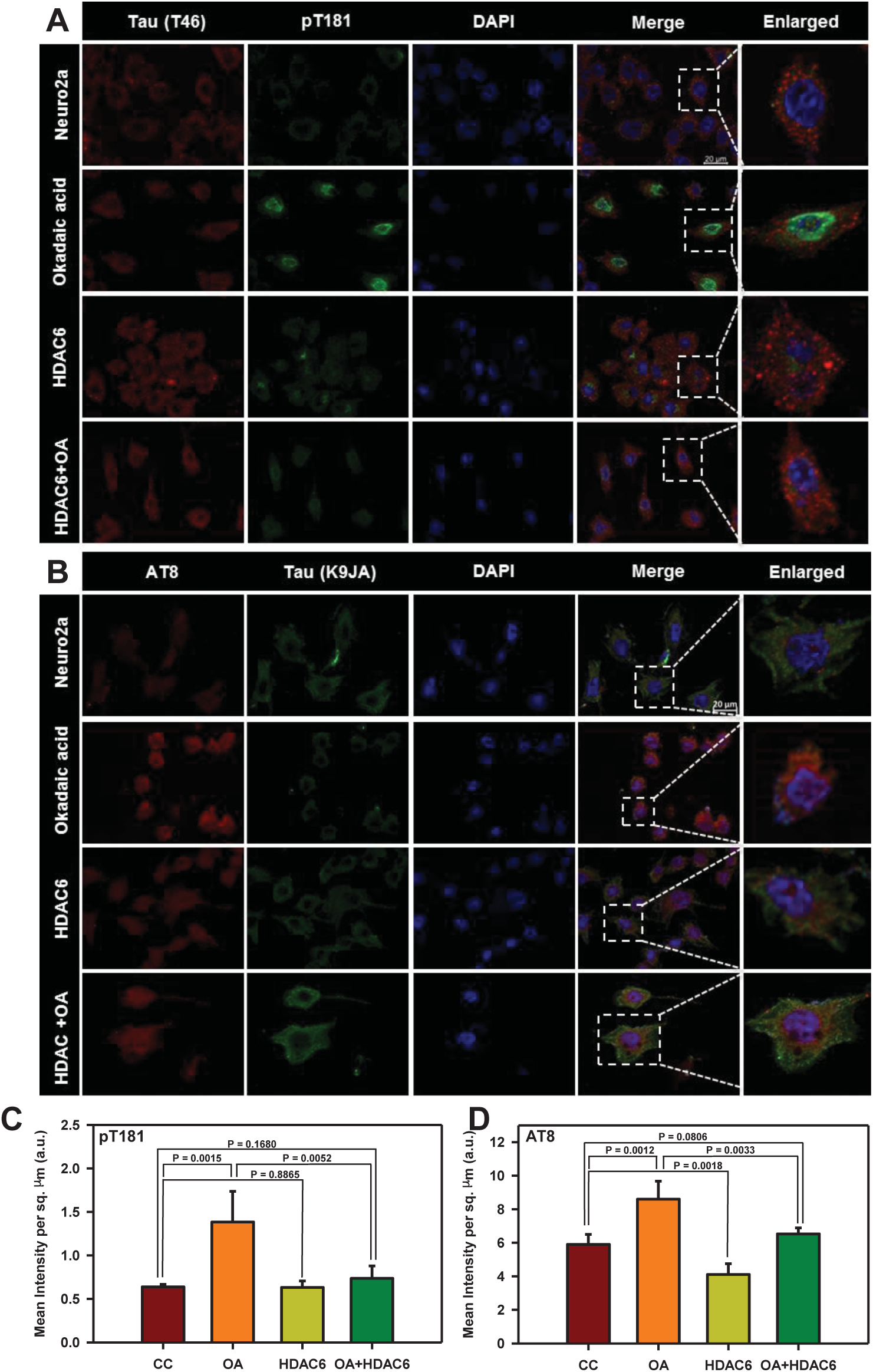
Inhibition of Tau phosphorylation by HDAC6. Neuro2a cells treated with okadaic acid (OA) enhanced overall phosphorylation by inhibition of PP2A. Tau phosphorylations at pT181 and AT8 epitopes of Tau were mapped after OA treatment alone as well as in simultaneous treatment with HDAC6 ZnF UBP to observe the effect on Tau phosphorylation. A) HDAC6 treatment for 24 hours along with OA shows inhibition of Tau phosphorylation at pT181 epitope whereas OA treatment shows increased phosphorylation at pT181. B) Phosphorylation at AT8 epitope is also reduced in presence of HDAC and OA as compared to OA alone. The enlarged image of neuro2a treated with OA alone and OA along with HDAC6 ZnF UBP shows marked difference in the level of phosphorylation at pT181 and AT8 epitopes. C, D). Mean fluorescence intensity for AT8 immunostaining of untreated neuro2a (CC), okadaic acid treatment (OA), HDAC6 ZnF UBP treatment (HDAC6) and HDAc6 ZnF UBP treatment along with okadaic acid. Okadaic acid treatment resulted in increased level of pT181 and AT8 phospho-epitopes. HDAC6 ZnF UBP treatment with OA reduced the level of both phospho-epitopes.

### HDAC6 ZnF UBP modulates actin dynamics

Actin organization and dynamics is crucial for cell shape and migration in respective cell types. In neuronal cells, actin dynamics is important for cell-to-cell communication through regulating neurite extension. Actin assembly and dynamics requires complex machinery and nucleating factors [46,47]. In neurodegenerative diseases, the regulation of actin dynamics gets impaired resulting in loss of synapses and dendritic spines [48]. HDAC6 is known to modulate actin organization through its association and deacetylase activity on cortactin and arp2/3 complex [24,46,47]. The function of HDAC6 ZnF UBP domain in actin dynamics is not yet understood. Our present observations showed HDAC6 ZnF UBP treatment lead to increase in neuritic extensions in neuro2a cells. We mapped HDAC6 ZnF UBP treated cells with FITC-Phalloidin to observe the F-actin cytoskeleton. We studied actin organization along with Tau and it was observed that HDAC6 ZnF UBP treatment had no effect on Tau levels but the actin cytoskeleton changed significantly (Fig.4A). Orthogonal sections were taken to clearly visualize the actin extensions. The initial orthogonal sections showed numerous extensions in HDAC6 ZnF UBP treated cells as compared to control (Fig.4B). As previously observed, HDAC6 ZnF UBP treated cells showed more neuritic extensions and dense actin cytoskeleton. Tau and actin are known to co-localize in neuritic extensions in previous studies where they facilitate growth cone transition [49]. We also observed Tau-actin co-localization in neuritic extensions indicating the interplaying of microtubule and actin dynamics in formation of cell extensions (Fig 4B, Supp. Fig. S2A, B). Greyscale images for actin immunostaining shows localization in neurite extensions and growth cones in filopodia-like structures (Fig. 4C, D).

**Figure 4.**
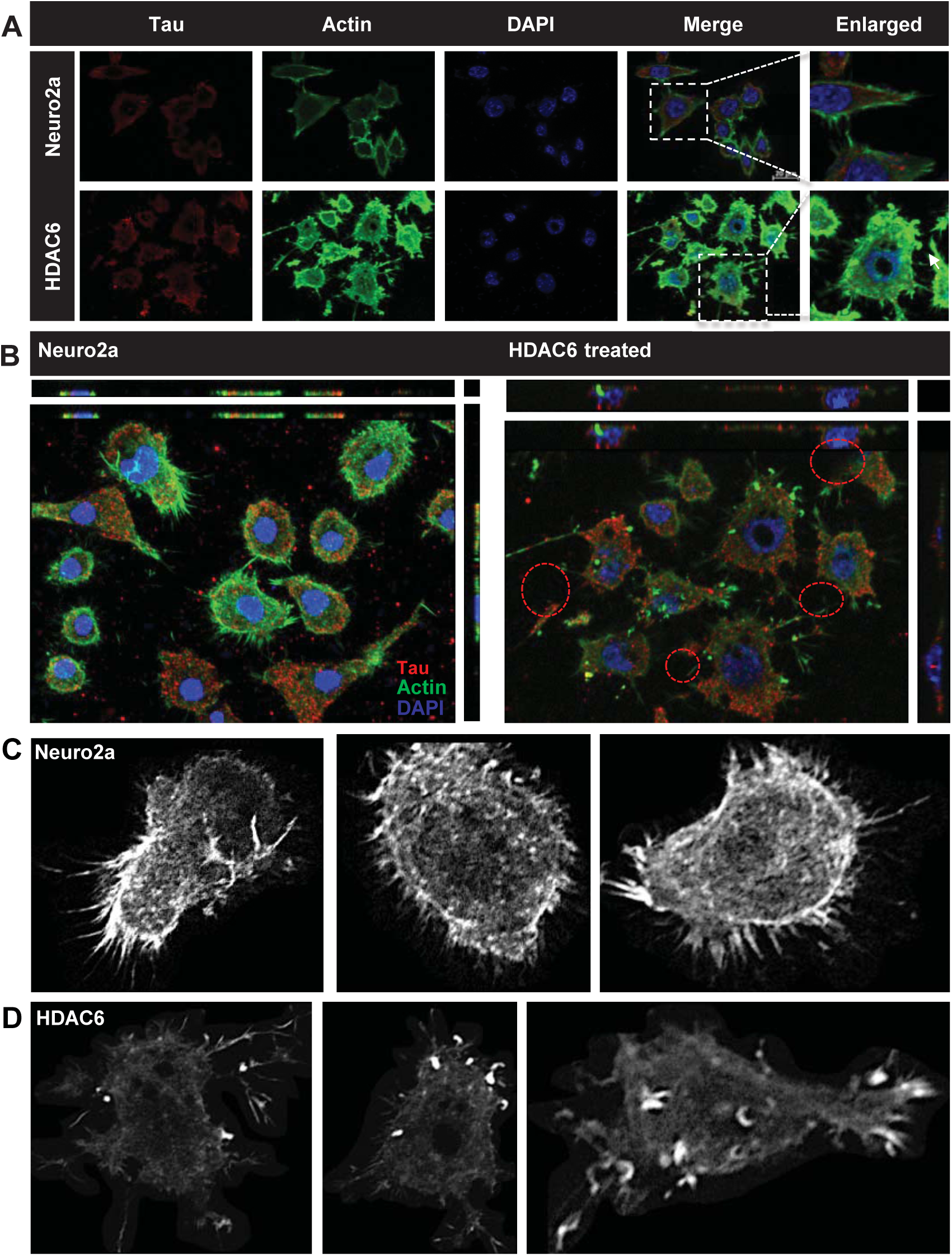
HDAC6 as a modulator of actin dynamics. HDAC6 deacetylase activity is known to modulate actin dynamics. The effect on actin dynamics was studied after exposure to HDAC6 ZnF UBP domain. HDAC6 ZnF UBP treatment to neuro2a resulted in enhancement of neurite extensions which are mapped by F-actin staining. A) HDAC6 ZnF UBP treatment for 24 hours increases the neuronal extensions as compared to untreated control cells when the extensions are mapped by actin immunostaining. The Tau levels remain unaltered in both the groups. B) The orthogonal projection shows the increase in number of neurites in the HDAC6 ZnF UBP treated cells. C) Greyscale images for untreated neuro2a cells mapped for F-actin showed small extensions in the growth cones. D) In HDAC6 ZnF UBP treated cells, F-actin containing longer extensions were observed. Growth cones were observed to be concentrated in membrane extensions and filopodia-like structures.

### Enhancement of podosome-like structures by HDAC6 ZnF UBP

Podosomes are actin based structures involved in the remodeling of extracellular matrix (ECM) and cell migration while podonuts consists of a cluster of podosomes interacting with ECM [50]. It consists of an actin rich core surrounded by actin regulatory molecules-like cortactin and arp2/3 as well as cell adhesion molecules like Talin and Vinculin [51]. Morphological changes in neuronal cells were observed when treated with HDAC6 ZnF UBP. HDAC6 ZnF UBP treated cells showed increased membrane ruffles and podosome-like structures (Supp. Fig. 3). Neuro2a cells were incubated with 50 nM HDAC6 ZnF UBP on 18 mm coverslips for 24 hours prior to immunostaining preparation. Immunostaining showed the localization of actin in the membrane ruffles in HDAC6 ZnF UBP treated cells. We observed minimal neuritic extensions in untreated neuro2a as compared to treated cells (Fig. 5A). Actin localization was observed mostly along the cell periphery in untreated neuro2a while it was focused in the neurite extensions and protruding podosome structure in HDAC6 ZnF UBP treated cells (Fig. 5B). Further, to check for HDAC6 localization on treatment, we mapped actin with HDAC6 ZnF UBP. The control cells showed a normal F-actin cytoskeleton in the extensions but HDAC6 treated cells showed presence of actin along with HDAC6 in these extensions (Fig.5C). Involvement of podosomes in cell adhesion and migration is an important attribute of invasive cells. Neuro2a, being a neuroblastoma cell line can form podosomes for cell migration and attachment. In contrast to untreated control neuronal cells (Fig.6A, B, C), HDAC6 ZnF UBP treated cells showed enhanced cell extensions and actin rich structures similar to podosomes formed in migrating and invading cells. HDAC6 treated cells showed a variety of actin-based membrane protrusions which can be morphologically classified as filopodia or lamellipodia, podosomes and podonuts (Fig. 6D-J). The neuritic extensions and podosomes were quantified in untreated control cells (CC) and HDAC6 treated cells. The neurites and podosomes were counted manually in multiple fields for untreated control and HDAC6 treated group (No. of fields, n = 6). HDAC6 ZnF UBP treated cells showed significantly enhanced neurite extensions and podosomes compared to untreated cells implicating its effect on actin dynamics and re-organization (Fig. 6K, L).

**Figure 5.**
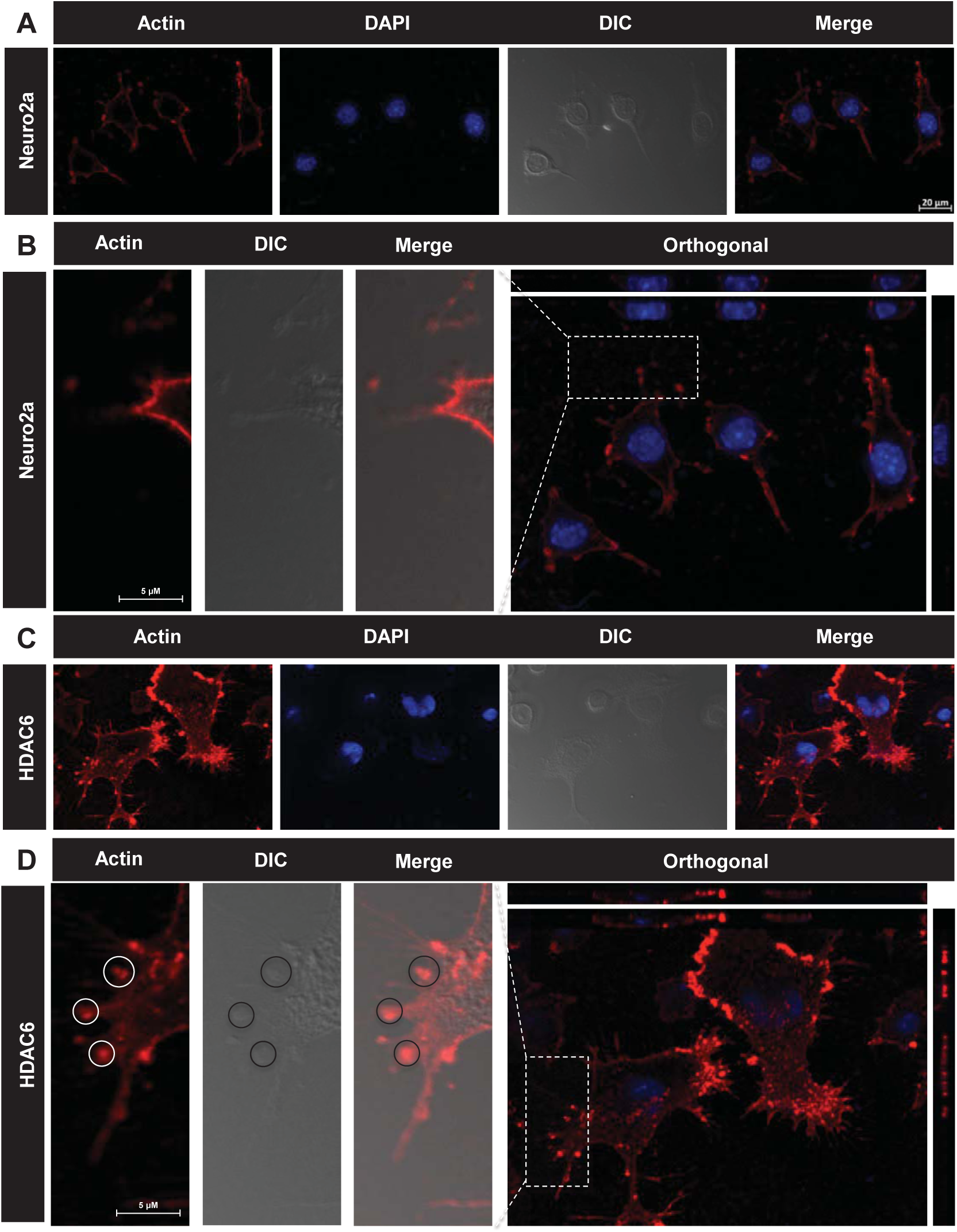
Enhancement of podosome formation by HDAC6. HDAC6 deacetylase activity is known for regulating the actin dynamics in cells and podosome formation. Structures resembling podosomes and podonuts were observed in neuro2a upon HDAC6 ZnF UBP treatment. Podosomes and podonuts are actin rich structures involved in cell attachment and migration. A) Podosomes marked by the actin rich structures along plasma membrane, were observed in neuro2a cells at a minimal level. B) Enhancement in podosome like structures observed in HDAC6 ZnF UBP treated cells. Orthogonal projection images showed marked difference in the membrane morphology and actin concentrated in membrane ruffles and podosome like structures in HDAC6 ZnF UBP treated cells. C) HDAC6 is found to be present in the neuritic extensions along with actin suggesting its role in regulation of neurite extensions. This is not seen in case of untreated cells as indicated in the enlarged images for both groups.

**Figure 6.**
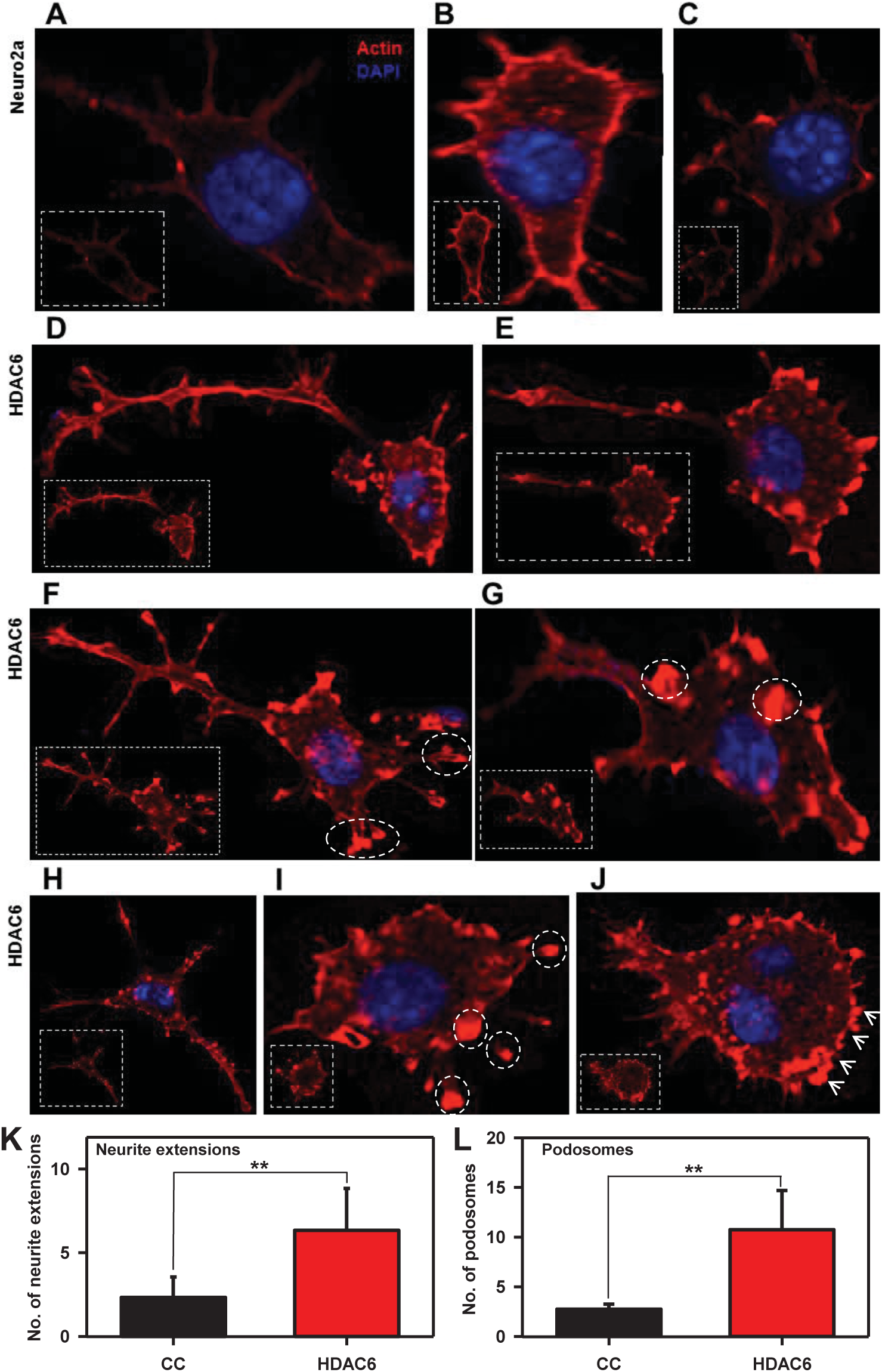
Podosome, lamellipodia and podonut-like structure induced by HDAC6 ZnF UBP. Neuro2a cells exposed to HDAC6 ZnF UBP exhibit a variety of actin rich structures characteristics of migratory or invading cells. A, B, C) Untreated neuro2a cells showed actin uniformly distributed along the periphery with small neuritic extensions while D, E) HDAC6 ZnF UBP treated cells showed longer extensions and membranes ruffles resembling podosomes involved in cell migration. F, G) Invadopodia and lamellipodia-like structures (encircled) were also observed, which are found in phagocytic cells. H, I, J) Most of the treated cells were observed to consist of assemblance of podosomes (encircled) and podosome clusters called podonuts (white arrow) rich in actin. K) The overall neurite extensions in neuro2a cells were counted in untreated and HDAC6 treated group in β-actin immunostained cells. The number of extensions in HDAC6 treated cells were found to be greatly enhanced compared to untreated group. L) The podosomes formed after HDAC6 ZnF UBP treatment were quantified by counting the podosome crown structures in both untreated and treated groups. HDAC6 ZnF UBP treated cells showed more podosome clusters signifying its actin modulating effect.

### HDAC6 ZnF UBP enhances ApoE nuclear localization

ApoE functions mainly as a lipid carrier in CNS where it delivers cholesterol to neurons *via* ApoE receptors following neuronal injury [52]. ApoE can be localized to nucleus and carries out induction of gene expression involved in inflammatory response. ApoE is the major apolipoprotein of central nervous system, produced primarily by astrocytes. ApoE is taken up by neurons where it plays important role in membrane maintenance and repair [53]. In normal physiological conditions, ApoE is localized in cytosol. ApoE nuclear localization has been reported in ovarian cancer cells, where it leads to better survival possibly through gene regulation [54]. We treated neuro2a cells with 50 nM of HDAC6 ZnF UBP domain to observe its effect on distribution and localization of ApoE in cell. It was observed that upon HDAC6 ZnF UBP treatment, there is increased nuclear localization of ApoE (Fig. 7A). The localization of ApoE was quantified by analyzing the intensity of ApoE immunofluorescence in nucleus and cytoplasm of untreated and HDAC6 ZnF UBP treated cells. The ApoE level in cytoplasm of both untreated and HDAC6 treated cells showed no change while it was significantly increased in the nucleus of HDAC6 ZnF UBP treated cells (Fig. 7B). The enhanced nuclear localization of ApoE may indicate in improved neuronal health [55]. ApoE can get translocated to the nucleus through a nuclear targeting chaperone nucleolin [54]. ApoE consists of weak nuclear localizing sequence, which indicates that there must be other mechanism involved in ApoE nuclear transport. Nucleolin is known to associate with apolipoproteins facilitating their translocation [56]. HDAC6 may mediate ApoE translocation through HSP90 as it is known to stabilize nucleolin during mitosis [57]. However, the exact mechanism needs to be further elucidated (Fig. 7C).

**Figure 7.**
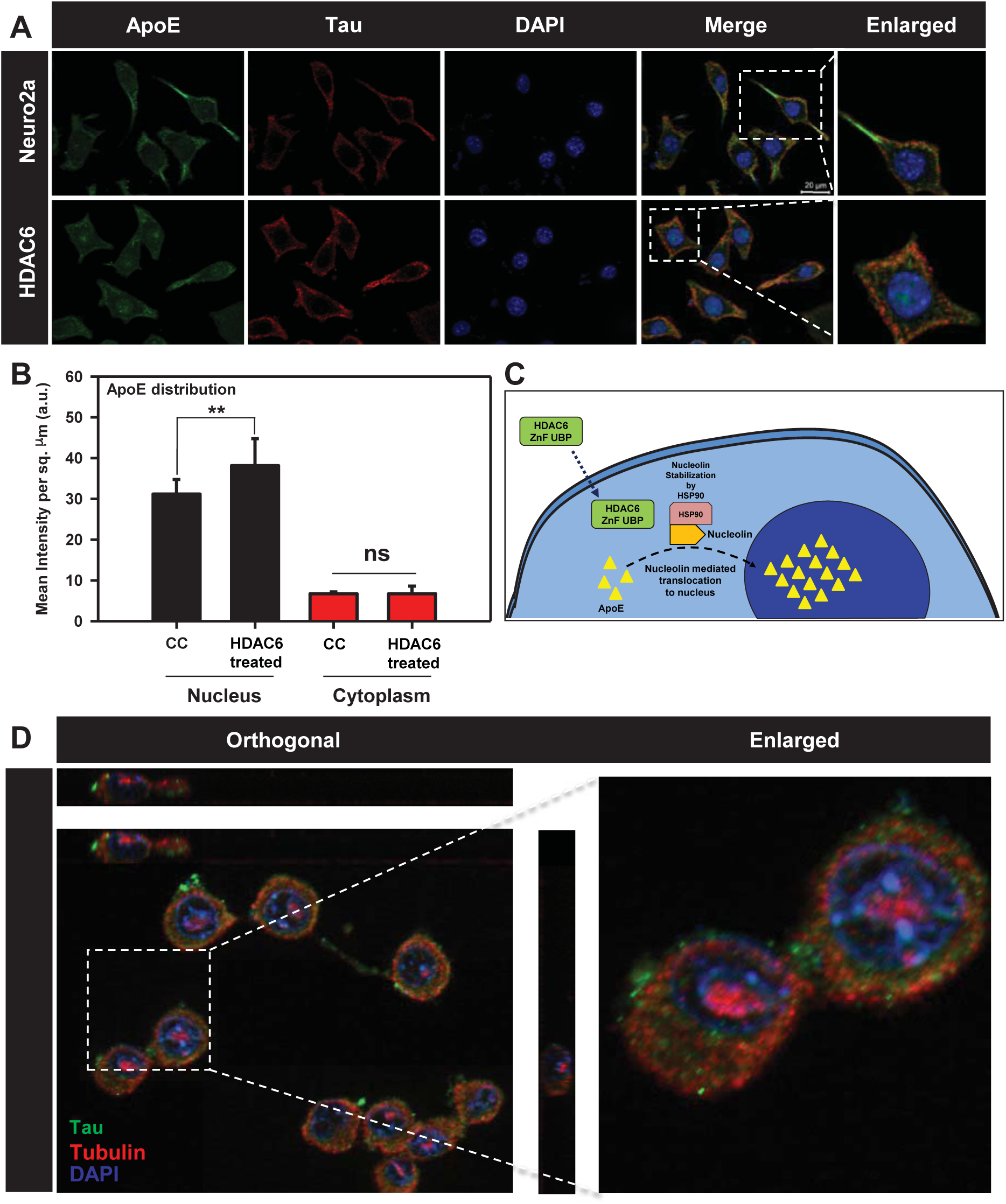
Modulation of ApoE and Tubulin localization by HDAC6. Cells were mapped for ApoE to observe its level and localization. Enhanced nuclear localization of ApoE was observed with HDAC6 ZnF UBP treatment. HDAC6 ZnF UBP also affected the tubulin distribution where the treated cells showed more tubulin localization in MTOC compared to untreated cells where tubulin was evenly distributed. A) Immunofluorescence mapping for ApoE and Tau upon HDAC6 treatment in neuro2a cells shows increased nuclear localization of ApoE. B) ApoE intensity in nucleus and cytoplasm was quantified for untreated and HDAC6 ZnF UBP treated neuro2a immunostained with anti-ApoE antibody. There was no significant difference in cytoplasmic level of ApoE in both groups while the nuclear ApoE fraction was notably increased in HDAC6 ZnF UBP treated cells. C) Enhanced ApoE localization upon HDAC6 ZnF UBP treatment may be mediated by nucleolin dependent transport of ApoE. ApoE requires an active transport system as it contains weak NLS signal. Nucleolin may serve this function as it is a known interacting partner of apolipoproteins and is stabilized by HSP90 whose activity is regulated by HDAC6. D) Neuro2a treated with HDAC6 ZnF UBP shows localization of tubulin focused over the nuclear periphery compared to untreated neuro2a where tubulin is evenly distributed in cytoplasm. Orthogonal projection image shows tubulin localization in nuclear periphery in HDAC6 ZnF UBP treated neuro2a cells.

### Enhanced Tubulin localization to MTOC with HDAC6 ZnF UBP treatment

[HDAC6 is a known interacting partner of tubulin and regulates microtubule structure and function through deacetylation. In a previous study, it was found that HDAC6 knockdown in cells along with HDAC6 inhibitor tubacin treatment does not affect microtubule growth velocity [58]. This implies that effect of HDAC6 independent of its deacetylase activity may also exist. We have given HDAC6 ZnF UBP to neuro2a cells to observe the effect on microtubule network. HDAC6 ZnF UBP treatment to neuro2a cells increased tubulin localization around nucleus in microtubule-organizing center (MTOC) (Fig. 7D). When neuro2a cells were mapped for actin and tubulin, untreated cells showed more axonal localization of tubulin along with actin, while HDAC6 treated cells showed tubulin predominantly in MTOC (Supp. Fig. 2A, B). The re-orientation of MTOC is a complex process which occurs in a dynein, cdc42 and dynactin dependent manner [59,60]. However, the mechanism and function of MTOC re-orientation is poorly understood. The results suggest the possible role of HDAC6 ZnF UBP domain in microtubule organization.

**Figure 8.**
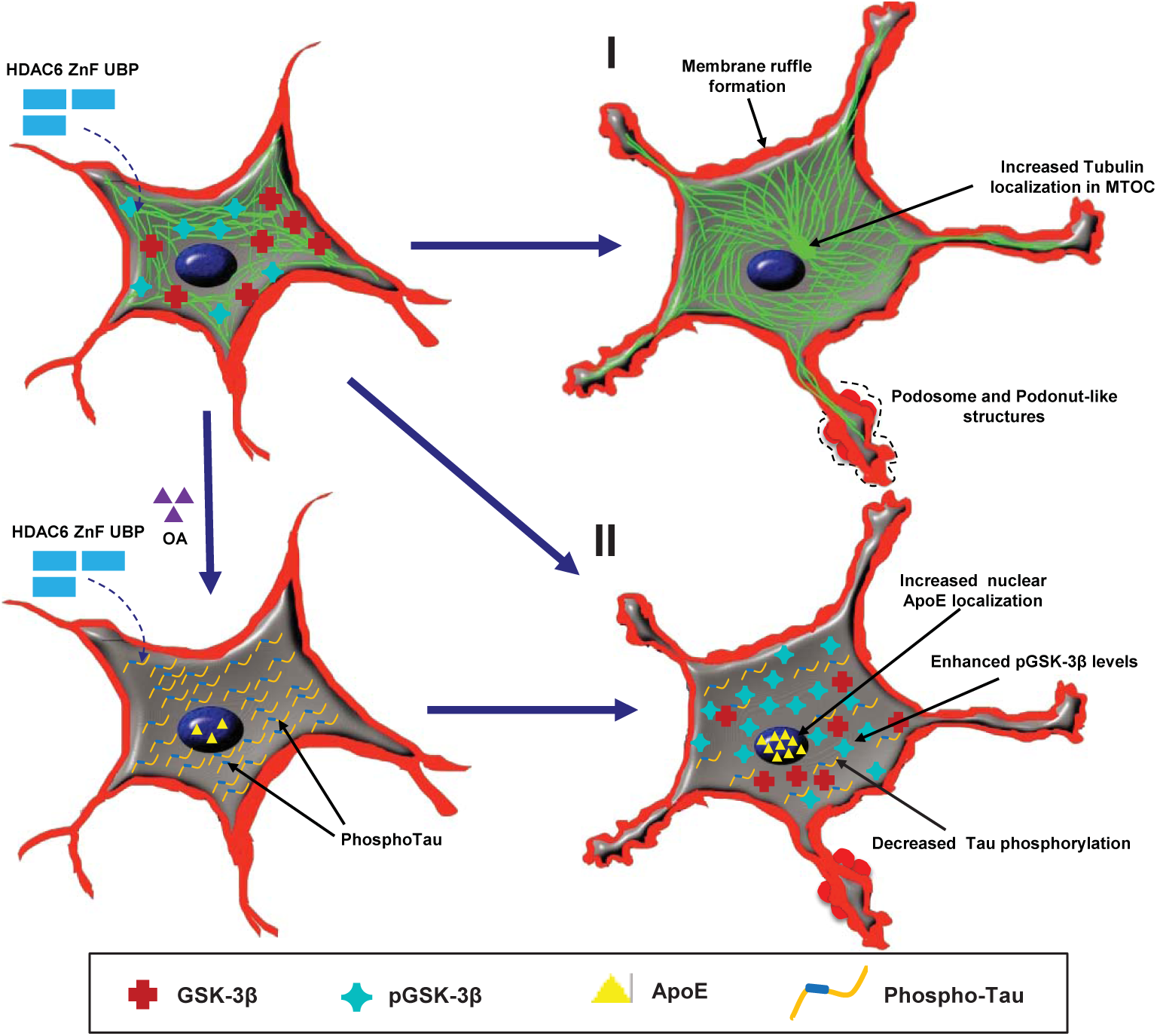
Effect of HDAC6 ZnF UBP on various cellular functions. HDAC6 ZnF UBP domain was observed to play possible roles in different aspects of neuronal cell function and morphology. Broadly classifying, we observed their effect in remodelling actin and tubulin as well as regulating Tau phosphorylation and GSK-3β activity. I) HDAC6 ZnF UBP treatment to the cells resulted in formation of podosome or lamellipodia-like structures and enhanced neurite extension. On the other hand, tubulin was found to more localized in MTOC in HDAC6 treated cells while untreated cells showed uniform tubulin distribution. II) The phosphorylation of Tau was also found to be reduced when HDAC6 ZnF UBP was given to the cells along with OA as inducer of phosphorylation. OA treated cells showed increased PhosphoTau levels (pT181 and AT8) while the levels of both were reduced in treated cells. The level of pGSK-3β was found to be increased with HDAC6 ZnF UBP treatment while total GSK-3β remains unaltered. Another aspect of HDAC6 treatment was observed in the localization of ApoE. Enhanced nuclear localization was found in treated cells suggesting the role of HDAC6 in regulating ApoE-mediated functions.

## DISCUSSION

Histone deactylases are known to be the enzymes that act on histones and regulates the epigenetic function in nucleus. However, many non-histone substrates of HDACs in the cytoplasm have been established [23]. Among the classes of histones deacetylase, class IIb deacetylase HDAC6 holds a distinctive position being the major cytoplasmic deacetylase having two active catalytic domains [23,61]. HDAC6 exhibit domainwise function, which may or may not be dependent on each other. HDAC6 plays a major role in cell proliferation, cell migration, misfolded protein degradation and stress response through its deacetylase function or interacting with various other proteins [62]. It is a well-known fact that poly-ubiquitinated protein aggregates are recruited through HDAC6 ZnF UBP to be sequestered into aggresomes. HDAC6 ZnF UBP binds to diglycine motif present on ubiquitin C-terminal, which gets exposed by deubiquitinase Ataxin3 [63]. The poly-ubiquitinated aggregates are carried *via* dynein motor to MTOC in order to form aggresomes. The sequestered aggregates in the form of aggresomes are cleared up by autophagy pathway [64,65]. HDAC6 levels are found to be elevated in the AD patients following increase in the aggregate burden in cells [29]. This suggests the possibility of involvement of this domain in tackling aggregates in a deacetylase independent manner. Neuro2a cells treated with various concentrations of HDAC6 ZnF UBP to assess their viability upon treatment using MTT assay and LDH assay. Neuro2a showed no toxicity in the presence of HDAC6 ZnF UBP. pT181 and AT8 (pS202/pT205) are two pathological phosphorylation events of Tau implicated in AD [30]. Okadaic acid treatment increases the level of these two epitopes by inhibition of PP2a activity. Treatment with HDAC6 ZnF UBP decreased the levels of both pT181 and AT8. The effect possibly arises from replenishing the PP2a activity upon inhibition by OA rather than inhibition of phosphorylation [41] Class II HDACs - HDAC1, HDAC6 and HDAC10 can form molecular complexes with phosphatases (PP1 or PP2a). However, HDAC6 was shown to interact only with the catalytic subunit of PP1 to form a complex retaining both phosphatase and deacetylase catalytic activity [66]. The binding of HDAC6 to PP1 was mapped to the second catalytic domain and C-terminal domain of HDAC6, which corresponds to ZnF UBP domain [41]. GSK-3β is a versatile protein kinase with more than hundred substrates [43]. GSK-3β acts on Tau in two different manners *i.e.* on pre-phosphorylated primed Tau and unprimed Tau [12]. In order for GSK-3β to work on primed substrate, the N-terminal primed substrate binding domain provides binding-site for primed substrates upregulating the kinase activity of GSK-3β. Phosphorylation at Serine 9 is the regulatory mechanism to down-regulate GSK-3β activity [11]. N-terminal domain with pSer9 acts as a pseudosubstrate for the primed substrate binding-domain of GSK-3β restricting the binding of other primed substrates like Tau [67]. HDAC6 ZnF UBP was found to increase the level of pSer9 phosphorylation on GSK-3β indicative of down-regulated GSK-3β activity with HDAC6 treatment in neuro2a cells.

Cellular functions like cell growth, morphogenesis, migration, intracellular transport and attachment require co-ordinated operation of cytoskeletal networks of actin and microtubules. Both Tau and HDAC6 are known to be involved in the regulation of both these cytoskeletal networks [68]. Although HDAC6 exerts its regulatory function through catalytic domain, but the role of HDAC6 ZnF UBP domain in this aspect need to be explored. We studied the effect of HDAC6 ZnF UBP, on cytoskeletal organization and found the ability of this domain to restructure cytoskeletal network. HDAC6 ZnF UBP treatment resulted in enhancement of neuritic extensions. HDAC6 is an important modulator of cell migration and structure by associating with actin and tubulin polymerization machinery of cells [24,58]. We observed the co-localization of actin and HDAC6 in the neurite extensions and formation of podosomes structures involved in cell migration in HDAC6 ZnF UBP treated cells. Podosomes are membranes invaginations formed in cells like macrophages, dendritic, smooth muscle, invasive cancer cells and in post-synaptic apparatus [69] The results showed the ability of HDAC6 ZnF UBP to enhance cell migration and neurite extensions by promoting the actin assembly along cell periphery. The role of HDAC6 in cytoskeletal organization has been studied extensively. HDAC6 has been reported to play regulatory role in chemotactic movement and migration of lymphocytes in deacetylase independent manner [70]. HDAC6 also regulate the membrane ruffle formation in HSP90 and Rac1 mediated mechanism and found to be localized in membrane ruffles along with actin [71]. In this aspect, the role of HDAC6 ZnF UBP in promoting neuritic extensions by actin remodelling can prove to be a novel aspect of HDAC6 in neuroprotection. HDAC6 ZnF UBP treatment also leads to enhanced localization of tubulin around nucleus signifying the possible structural and pathological aspect of HDAC6 ZnF UBP. The function of HDAC6 ZnF UBP domain is known with respect to aggregate clearance by mediating aggresome formation. However, current findings suggest that the domain may have other regulatory functions. Overall, our studies suggest the role of HDAC6 ZnF UBP domain in modulating Tau protein phosphorylation and its cytoskeletal organization which are independent of HDAC6 catalytic activity.

### Summary statement

HDAC6 ZnF UBP modulates actin organization to direct formation of podosome-like structure and regulates phosphorylation in neuronal cells.

## Materials and Methods

### Chemicals and reagents

Luria-Bertani broth (Himedia); Ampicillin, NaCl, Phenylmethylsulfonylfluoride (PMSF), MgCl_2_, APS, DMSO, Ethanol (Mol Bio grade), Isopropanol (Mol Bio grade) were purchased from MP biomedicals; IPTG and Dithiothreitol (DTT) from Calbiochem; MES, BES, SDS from Sigma; EGTA, Protease inhibitor cocktail, Tris base, 40% Acrylamide, TEMED from Invitrogen. For cell culture studies, Dulbecco modified eagle’s media (DMEM), Fetal bovine Serum (FBS), Horse serum, Phosphate buffer saline (PBS, cell biology grade), Trypsin-EDTA, Penicillin-streptomycin, Pierce™ LDH Cytotoxicity Assay Kit (Thermo, cat no 88953), RIPA buffer were also purchased from Invitrogen. MTT reagent, Okadaic acid and TritonX-100 were purchased from Sigma. The coverslip of 0.17 mm was purchased from Bluestar for immunofluorescence. In immunofluorescence and western blot study we used the following antibodies: Beta-actin (Thermofisher cat no. MA515739) Beta Tubulin (BT7R) (Thermofisher, cat no MA516308) and total Tau antibody K9JA (Dako, cat no A0024), pT181 (Invitrogen, cat no 701530) AT8 (Thermo fisher, cat no MN1020), GSK-3β (Thermo fisher, cat no MA5-15109), Phospho-GSK-3β (Ser9) (Thermo fisher, cat no MA5-14873), anti-ApoE (Sigma, cat no. SAB2701946), anti-mouse secondary antibody conjugated with Alexa Fluor-488 (Invitrogen, cat no A-11001), Goat anti-Rabbit IgG (H+L) Cross-Adsorbed Secondary Antibody with Alexa Fluor 555 (A-21428), Rabbit anti-Goat IgG (H+L) Cross-Adsorbed Secondary Antibody with Alexa Fluor 594 (A27016) and DAPI (Invitrogen).

### Protein expression and Purification

Full length Tau (hTau40wt) in pT7C were transformed and expressed in BL21* cells while HDAC6 in pET28a-LIC was transformed and expressed in BL21 Codon plus RIL cells. Full-length Tau and repeat domain Tau were purified in two steps using cation exchange chromatography and Size-exclusion chromatography. The cells expressing these proteins after transformation were scaled up and harvested. The cells were lysed by homogenization at 15000 KPSI. The lysate was supplied with 0.5 M NaCl and 5 mM DTT and kept at 90°C for 20 minutes to denature all the structured protein. The resulting sample was centrifuged at 40000 rpm for 45 minutes. The supernatant was kept for overnight dialysis in 20mM MES pH 6.8. The dialyzed sample was centrifuged again at 40000 rpm for 45 minutes and the supernatant was filtered and loaded onto Sepharose fast flow (SPFF) column pre-equilibrated with 20 mM MES pH 6.8, 50 mM NaCl. Elution was carried out using 20 mM MES pH 6.8, 1 M NaCl. The fractions collected from cation exchange chromatography containing Tau protein were pooled, concentrated and subjected to Size-exclusion chromatography using 1X PBS, 2 mM DTT in Superdex 75 Hi-load 16/600 column [72,73]. Purification of HDAC6 ZnF UBP was carried out by Ni-NTA affinity chromatography using 50 mM Tris-Cl pH 8.0 with 20 mM Imidazole for wash and 1000 mM imidazole for elution. The sample was dialyzed overnight in 50 mM Tris-Cl pH 8.0, 100 mM NaCl, 2.5 % glycerol to remove imidazole followed by Size-exclusion chromatography using Superdex 75 Hi-load 16/600 column [63].

### Cell viability by MTT assay

The effective concentration of HDAC6 for the subsequent treatments was determined by studying the concentration dependent toxicity studies by MTT assay. 10^4^ neuro2a cells were seeded in a 96 well culture plate in DMEM supplemented with 10% FBS and antibiotic penicillin-streptomycin for 24 hours at 37 °C CO_2_ incubator. The cells were treated with HDAC6 (0-500 nM) in serum-starved media for 24 hours. MTT at the concentration of 0.5 mg/mL was added to the cells and incubated for 3 hours. The reduction of MTT by cellular enzymes forms formazon crystals, which were dissolved in DMSO, and the colour developed was quantified by reading at 570 nm in a TECAN Infinite 200 PRO plate reader.

### LDH assay

The effect of HDAC6 treatment on cell membrane integrity was studied by LDH (Lactate Dehydrogenase) assay. The disruption of cell membrane integrity leads to the leakage of LDH enzyme, which is quantified by an enzymatic reaction giving a colored end product. For performing LDH assay the cells were incubated and treated as mentioned for MTT assay. After the treatment with HDAC 6 supernatant media was used and the assay was performed according to the manufacturer’s protocol. In brief, 50 µL of cell supernatant was incubated with 50 µL of the reaction mixture provided for 30 minutes at room temperature. 50 µL of stop solution was added to each well and the colour developed was measured at 490 nm and background subtraction at 680 nm was done.

### Caspase 3/7 activity assay

In order to study the effect of HDAC6 on inducing apoptotic cell death, the activity of executioner caspase 3 was determined by EnzChek™ Caspase-3 Assay Kit. 10000 cells/well cells were seeded in a 12 well culture plate for 24 hours and further treated with HDAC6 (0-500 nM) for 24 hours in serum-starved media. Caspase activity was performed as per manufacturer’s protocol. The cells were lysed with provided lysis buffer in freeze-thaw cycles. The cell debris was centrifuged out and the supernatant was incubated with fluorescent substrate (DEVD-Rhodamine). The fluorescence as quantified at (Ex/Em) 496/520 nm at different time intervals in TECAN Infinite 200 PRO plate reader.

### Cell culture and Immunofluorescence

Neuro2a cells were cultured in advanced DMEM supplemented with penstrep-glutamine, anti-mycotic and 10% FBS. For immunofluorescence studies 5×10^4^ cells were seeded on a glass coverslip (Bluestar) in a 12 well culture plate. Cells were given the desired treatment in serum starved media (0.5% FBS) for 24 hours including groups involving Okadaic acid (OA) treatment. After incubation period, cells were washed with PBS and fixed with 4% paraformaldehyde. Further cells were washed with PBS thrice and permiabilized using 0.2% Triton X-100. Cells were blocked with 2% horse serum and incubated with primary antibodies in a moist chamber at 4 ºC overnight. Next day, cells were washed thrice with 1X PBS and incubated with alexa fluor labeled secondary antibodies for 1 hour at 37 ºC. The unbound secondary antibody was washed off with three washes of PBS and counterstained with DAPI. The coverslips were mounted in 80% glycerol and observed under 63X oil immersion lens in Axio Observer 7.0 Apotome 2.0 (Zeiss) microscope.

### Statistical analysis

Two-tailed unpaired student t-test was used to determine the significance for experiments involving comparison of two groups (n.s. – non-significant, * indicates P ≤0.05, ** indicates P ≤ 0.01, *** indicates P ≤ 0.001). One way ANOVA was conducted for the levels of phospho-epitopes (pT181 and AT8) to compare different treatment groups. Tukey’s HSD (Honest significant difference) test was performed to compare the significance within groups (Significant at Tukey’s HSD p<0.05). All the experiments were performed in triplicates and analyzed by Sigmaplot 10.0 (Systat software).

## Author Contributions

AB, SKS and SC conducted most of the experiments, analyzed the results, and wrote the paper. SC conceived the idea for the project and wrote the paper.

## Notes

The authors have declared no conflict of interest with the contents of this article.

## ACKNOWLEDGEMENTS

This project is supported in part by grants from the Department of Biotechnology from Neuroscience Task Force (Medical Biotechnology-Human Development & Disease Biology (DBT-HDDB))-BT/PR/19562/MED/122/13/2016 and in-house CSIR-National Chemical Laboratory grant MLP029526. Abhishek Ankur Balmik acknowledges the Shyama Prasad Mukherjee fellowship (SPMF) from Council of Scientific Industrial Research (CSIR), India. SKS acknowledges Department of Biotechnology for fellowship. Tau constructs were kindly gifted by Prof. Roland Brandt from University of Osnabruck, Germany. HDAC6 ZnF UBP construct was kindly gifted by Prof. Cheryl Aerosmith, University of Toronto, Canada.

## Author details

**Abhishek A Balmik**

Neurobiology Group, Division of Biochemical Sciences, CSIR-National Chemical Laboratory, Dr. Homi Bhabha Road, 411008 Pune, India

Academy of Scientific and Innovative Research (AcSIR), 411008 Pune, India

**Contribution:** Methodology, Investigation, Validation, Formal analysis, Writing – original draft.

**Competing interests:** No competing interests declared.

**Shweta K Sonawane**

Neurobiology Group, Division of Biochemical Sciences, CSIR-National Chemical Laboratory, Dr. Homi Bhabha Road, 411008 Pune, India

Academy of Scientific and Innovative Research (AcSIR), 411008 Pune, India

**Contribution:** Methodology, Investigation, Validation, Formal analysis, Writing – original draft.

**Competing interests:** No competing interests declared.

**Subashchandrabose Chinnathambi**

Neurobiology Group, Division of Biochemical Sciences, CSIR-National Chemical Laboratory, Dr. Homi Bhabha Road, 411008 Pune, India

Academy of Scientific and Innovative Research (AcSIR), 411008 Pune, India

**Contribution:** Conceptualization, Resources, Supervision, Funding acquisition, Validation, Project administration, Writing—review and editing.

**Competing interests:** No competing interests declared.

## References

[1] Williams, D. (2006). Tauopathies: classification and clinical update on neurodegenerative diseases associated with microtubule-associated protein tau. Internal medicine journal 36, 652–660.

[2] Coppola, G. et al. (2012). Evidence for a role of the rare p. A152T variant in MAPT in increasing the risk for FTD-spectrum and Alzheimer’s diseases. Human molecular genetics 21, 3500–3512.

[3] Ksiezak-Reding, H. and Yen, S.-H. (1991). Structural stability of paired helical filaments requires microtubule-binding domains of tau: a model for self-association. Neuron 6, 717–728.

[4] Iqbal, K., Liu, F., Gong, C.-X. and Grundke-Iqbal, I. (2010). Tau in Alzheimer disease and related tauopathies. Current Alzheimer Research 7, 656–664.

[5] Goedert, M., Ghetti, B. and Spillantini, M.G. (2012). Frontotemporal dementia: implications for understanding Alzheimer disease. Cold Spring Harbor perspectives in medicine 2, a006254.

[6] Martin, L., Latypova, X. and Terro, F. (2011). Post-translational modifications of tau protein: implications for Alzheimer’s disease. Neurochemistry international 58, 458–471.

[7] Biernat, J., Gustke, N., Drewes, G. and Mandelkow, E. (1993). Phosphorylation of Ser262 strongly reduces binding of tau to microtubules: distinction between PHF-like immunoreactivity and microtubule binding. Neuron 11, 153–163.

[8] Biernat, J. et al. (1992). The switch of tau protein to an Alzheimer-like state includes the phosphorylation of two serine-proline motifs upstream of the microtubule binding region. The EMBO journal 11, 1593–1597.

[9] Drewes, G., Ebneth, A., Preuss, U., Mandelkow, E.-M. and Mandelkow, E. (1997). MARK, a novel family of protein kinases that phosphorylate microtubule-associated proteins and trigger microtubule disruption. Cell 89, 297–308.

[10] Lucas, J.J., Hernández, F., Gómez-Ramos, P., Morán, M.A., Hen, R. and Avila, J. (2001). Decreased nuclear β-catenin, tau hyperphosphorylation and neurodegeneration in GSK-3β conditional transgenic mice. The EMBO journal 20, 27–39.

[11] Cho, J.H. and Johnson, G.V. (2004). Primed phosphorylation of tau at Thr231 by glycogen synthase kinase 3β (GSK3β) plays a critical role in regulating tau’s ability to bind and stabilize microtubules. Journal of neurochemistry 88, 349–358.

[12] Cho, J.-H. and Johnson, G.V. (2003). Glycogen Synthase Kinase 3β Phosphorylates tau at both primed and unprimed sites differential impact on microtubule binding. Journal of Biological Chemistry 278, 187–193.

[13] Mandelkow, E.-M. and Mandelkow, E. (2012). Biochemistry and cell biology of tau protein in neurofibrillary degeneration. Cold Spring Harbor perspectives in medicine 2, a006247.

[14] Amm, I., Sommer, T. and Wolf, D.H. (2014). Protein quality control and elimination of protein waste: the role of the ubiquitin–proteasome system. Biochimica et Biophysica Acta (BBA)-Molecular Cell Research 1843, 182–196.

[15] Pierre, S.-R., Vernace, V., Wang, Z. and Figueiredo-Pereira, M.E. (2013) Assembly of protein aggregates in neurodegeneration: mechanisms linking the ubiquitin/proteasome pathway and chaperones. In Madame Curie Bioscience Database [Internet] ed.^eds). Landes Bioscience

[16] Ciechanover, A. and Kwon, Y.T. (2015). Degradation of misfolded proteins in neurodegenerative diseases: therapeutic targets and strategies. Experimental & molecular medicine 47, e147.

[17] Gorantla, N.V. and Chinnathambi, S. (2018). Tau Protein Squired by Molecular Chaperones During Alzheimer’s Disease. Journal of Molecular Neuroscience 66, 356–368.

[18] Tai, H.-C. and Schuman, E.M. (2008). Ubiquitin, the proteasome and protein degradation in neuronal function and dysfunction. Nature Reviews Neuroscience 9, 826.

[19] Casadei, N. et al. (2013). Overexpression of synphilin-1 promotes clearance of soluble and misfolded alpha-synuclein without restoring the motor phenotype in aged A30P transgenic mice. Human molecular genetics 23, 767–781.

[20] Ardley, H.C., Scott, G.B., Rose, S.A., Tan, N.G. and Robinson, P.A. (2004). UCH-L1 aggresome formation in response to proteasome impairment indicates a role in inclusion formation in Parkinson’s disease. Journal of neurochemistry 90, 379–391.

[21] Simões-Pires, C., Zwick, V., Nurisso, A., Schenker, E., Carrupt, P.-A. and Cuendet, M. (2013). HDAC6 as a target for neurodegenerative diseases: what makes it different from the other HDACs? Molecular neurodegeneration 8, 7.

[22] Verdin, E., Dequiedt, F. and Kasler, H.G. (2003). Class II histone deacetylases: versatile regulators. TRENDS in Genetics 19, 286–293.

[23] Yao, Y.-L. and Yang, W.-M. (2010). Beyond histone and deacetylase: an overview of cytoplasmic histone deacetylases and their nonhistone substrates. BioMed Research International 2011

[24] Zhang, X. et al. (2007). HDAC6 modulates cell motility by altering the acetylation level of cortactin. Molecular cell 27, 197–213.

[25] Pandey, U.B., Batlevi, Y., Baehrecke, E.H. and Taylor, J.P. (2007). HDAC6 at the intersection of autophagy, the ubiquitin-proteasome system, and neurodegeneration. Autophagy 3, 643–645.

[26] Pandey, U.B. et al. (2007). HDAC6 rescues neurodegeneration and provides an essential link between autophagy and the UPS. Nature 447, 860.

[27] Hao, R., Nanduri, P., Rao, Y., Panichelli, R.S., Ito, A., Yoshida, M. and Yao, T.-P. (2013). Proteasomes activate aggresome disassembly and clearance by producing unanchored ubiquitin chains. Molecular cell 51, 819–828.

[28] Iwata, A., Riley, B.E., Johnston, J.A. and Kopito, R.R. (2005). HDAC6 and microtubules are required for autophagic degradation of aggregated huntingtin. Journal of Biological Chemistry 280, 40282–40292.

[29] Zhang, L., Sheng, S. and Qin, C. (2013). The role of HDAC6 in Alzheimer’s disease. Journal of Alzheimer’s Disease 33, 283–295.

[30] Šimić, G. et al. (2016). Tau protein hyperphosphorylation and aggregation in Alzheimer’s disease and other tauopathies, and possible neuroprotective strategies. Biomolecules 6, 6.

[31] Šimić, G., Diana, A. and Hof, P.R. (2003) Phosphorylation pattern of tau associated with distinct changes of the growth cone cytoskeleton. In Guidance Cues in the Developing Brain ed.^eds), pp. 33–48. Springer

[32] Arendt, T., Holzer, M., Fruth, R., Brückner, M. and Gärtner, U. (1995). Paired helical filament-like phosphorylation of tau, deposition of β/A4-amyloid and memory impairment in rat induced by chronic inhibition of phosphatase 1 and 2A. Neuroscience 69, 691–698.

[33] Sontag, E., Hladik, C., Montgomery, L., Luangpirom, A., Mudrak, I., Ogris, E. and White III, C.L. (2004). Downregulation of protein phosphatase 2A carboxyl methylation and methyltransferase may contribute to Alzheimer disease pathogenesis. Journal of Neuropathology & Experimental Neurology 63, 1080–1091.

[34] Vintém, A.P.B., Henriques, A.G., e Silva, O.A.d.C. and e Silva, E.F.d.C. (2009). PP1 inhibition by Aβ peptide as a potential pathological mechanism in Alzheimer’s disease. Neurotoxicology and teratology 31, 85–88.

[35] Li, B., Chohan, M.O., Grundke-Iqbal, I. and Iqbal, K. (2007). Disruption of microtubule network by Alzheimer abnormally hyperphosphorylated tau. Acta neuropathologica 113, 501–511.

[36] Penzes, P. and VanLeeuwen, J.-E. (2011). Impaired regulation of synaptic actin cytoskeleton in Alzheimer’s disease. Brain research reviews 67, 184–192.

[37] Fiala, J.C., Spacek, J. and Harris, K.M. (2002). Dendritic spine pathology: cause or consequence of neurological disorders? Brain research reviews 39, 29–54.

[38] Selkoe, D.J. (2002). Alzheimer’s disease is a synaptic failure. Science 298, 789–791.

[39] Boyault, C., Sadoul, K., Pabion, M. and Khochbin, S. (2007). HDAC6, at the crossroads between cytoskeleton and cell signaling by acetylation and ubiquitination. Oncogene 26, 5468.

[40] Alao, J.P., Stavropoulou, A.V., Lam, E.W. and Coombes, R.C. (2006). Role of glycogen synthase kinase 3 beta (GSK3β) in mediating the cytotoxic effects of the histone deacetylase inhibitor trichostatin A (TSA) in MCF-7 breast cancer cells. Molecular cancer 5, 40.

[41] Brush, M.H., Guardiola, A., Connor, J.H., Yao, T.-P. and Shenolikar, S. (2004). Deactylase inhibitors disrupt cellular complexes containing protein phosphatases and deacetylases. Journal of Biological Chemistry 279, 7685–7691.

[42] Beurel, E. (2011). HDAC6 regulates LPS-tolerance in astrocytes. PLoS One 6, e25804.

[43] Beurel, E., Grieco, S.F. and Jope, R.S. (2015). Glycogen synthase kinase-3 (GSK3): regulation, actions, and diseases. Pharmacology & therapeutics 148, 114–131.

[44] Avila, J. (2008). Tau kinases and phosphatases. Journal of cellular and molecular medicine 12, 258–259.

[45] Boban, M., Leko, M.B., Miškić, T., Hof, P.R. and Šimić, G. (2018). Human neuroblastoma SH-SY5Y cells treated with okadaic acid express phosphorylated high molecular weight tau-immunoreactive protein species. Journal of neuroscience methods

[46] Oda, T., Iwasa, M., Aihara, T., Maéda, Y. and Narita, A. (2009). The nature of the globular-to fibrous-actin transition. Nature 457, 441.

[47] Lee, S.H. and Dominguez, R. (2010). Regulation of actin cytoskeleton dynamics in cells. Molecules and cells 29, 311–325.

[48] Moghaddam, H.S. and Aarabi, M.H. (2018). Aβ-Mediated Dysregulation of F-Actin Nanoarchitecture Leads to Loss of Dendritic Spines and Alzheimer’s Disease-Related Cognitive Impairments. Journal of Neuroscience 38, 5840–5842.

[49] Biswas, S. and Kalil, K. (2018). The microtubule-associated protein tau mediates the organization of microtubules and their dynamic exploration of actin-rich lamellipodia and filopodia of cortical growth cones. Journal of Neuroscience 38, 291–307.

[50] Gimona, M., Buccione, R., Courtneidge, S.A. and Linder, S. (2008). Assembly and biological role of podosomes and invadopodia. Current opinion in cell biology 20, 235–241.

[51] Siddiqui, T.A., Lively, S., Vincent, C. and Schlichter, L.C. (2012). Regulation of podosome formation, microglial migration and invasion by Ca 2+-signaling molecules expressed in podosomes. Journal of neuroinflammation 9, 250.

[52] Sen, A., Nelson, T.J. and Alkon, D.L. (2015). ApoE4 and Aβ oligomers reduce BDNF expression via HDAC nuclear translocation. Journal of Neuroscience 35, 7538–7551.

[53] Jiang, Q. et al. (2008). ApoE promotes the proteolytic degradation of Aβ. Neuron 58, 681–693.

[54] Kim, W.S., Elliott, D.A., Kockx, M., Kritharides, L., Rye, K.-A., Jans, D.A. and Garner, B. (2008). Analysis of apolipoprotein E nuclear localization using green fluorescent protein and biotinylation approaches. Biochemical Journal 409, 701–709.

[55] Rohn, T.T. and Moore, Z.D. (2017). Nuclear localization of apolipoprotein E4: a new trick for an old protein. International journal of neurology and neurotherapy 4

[56] Semenkovich, C.F., Ostlund Jr, R.E., Olson, M.O. and Yang, J.W. (1990). A protein partially expressed on the surface of HepG2 cells that binds lipoproteins specifically is nucleolin. Biochemistry 29, 9708–9713.

[57] Wang, S.-A., Li, H.-Y., Hsu, T.-I., Chen, S.-H., Wu, C.-J., Chang, W.-C. and Hung, J.-J. (2011). Heat shock protein 90 stabilizes nucleolin to increase mRNA stability in mitosis. Journal of Biological Chemistry 286, 43816–43829.

[58] Zilberman, Y., Ballestrem, C., Carramusa, L., Mazitschek, R., Khochbin, S. and Bershadsky, A. (2009). Regulation of microtubule dynamics by inhibition of the tubulin deacetylase HDAC6. Journal of cell science 122, 3531–3541.

[59] Sancho, D., Vicente-Manzanares, M., Mittelbrunn, M., Montoya, M.C., Gordón-Alonso, M., Serrador, J.M. and Sánchez-Madrid, F. (2002). Regulation of microtubule-organizing center orientation and actomyosin cytoskeleton rearrangement during immune interactions. Immunological reviews 189, 84–97.

[60] Palazzo, A.F., Joseph, H.L., Chen, Y.-J., Dujardin, D.L., Alberts, A.S., Pfister, K.K., Vallee, R.B. and Gundersen, G.G. (2001). Cdc42, dynein, and dynactin regulate MTOC reorientation independent of Rho-regulated microtubule stabilization. Current Biology 11, 1536–1541.

[61] Li, M., Zhuang, Y. and Shan, B. (2016) Analysis of expression and functions of histone deacetylase 6 (hdac6). In Histone Deacetylases ed.^eds), pp. 85–94. Springer

[62] Li, Y., Shin, D. and Kwon, S.H. (2013). Histone deacetylase 6 plays a role as a distinct regulator of diverse cellular processes. The FEBS journal 280, 775–793.

[63] Ouyang, H. et al. (2012). Protein aggregates are recruited to aggresome by histone deacetylase 6 via unanchored ubiquitin C termini. Journal of Biological Chemistry 287, 2317–2327.

[64] Yan, J. (2014). Interplay between HDAC6 and its interacting partners: essential roles in the aggresome-autophagy pathway and neurodegenerative diseases. DNA and cell biology 33, 567–580.

[65] Van Helleputte, L., Benoy, V. and Van Den Bosch, L. (2014). The role of histone deacetylase 6 (HDAC6) in neurodegeneration. Res Rep Biol 5, 1–13.

[66] Esteves, S.L., Domingues, S.C., da Cruz e Silva, O.A., Fardilha, M. and da Cruz e Silva, E.F. (2012). Protein phosphatase 1α interacting proteins in the human brain. Omics: a journal of integrative biology 16, 3–17.

[67] Fang, X., Yu, S.X., Lu, Y., Bast, R.C., Woodgett, J.R. and Mills, G.B. (2000). Phosphorylation and inactivation of glycogen synthase kinase 3 by protein kinase A. Proceedings of the National Academy of Sciences 97, 11960–11965.

[68] Elie, A. et al. (2015). Tau co-organizes dynamic microtubule and actin networks. Scientific reports 5, 9964.

[69] Vincent, C., Siddiqui, T.A. and Schlichter, L.C. (2012). Podosomes in migrating microglia: components and matrix degradation. Journal of neuroinflammation 9, 190.

[70] Cabrero, J.R. et al. (2006). Lymphocyte chemotaxis is regulated by histone deacetylase 6, independently of its deacetylase activity. Molecular biology of the cell 17, 3435–3445.

[71] Gao, Y.-s., Hubbert, C.C., Lu, J., Lee, Y.-S., Lee, J.-Y. and Yao, T.-P. (2007). Histone deacetylase 6 regulates growth factor-induced actin remodeling and endocytosis. Molecular and cellular biology 27, 8637–8647.

[72] Gorantla, N.V., Khandelwal, P., Poddar, P. and Chinnathambi, S. (2017) Global conformation of tau protein mapped by Raman spectroscopy. In Tau Protein ed.^eds), pp. 21–31. Springer

[73] Gorantla, N.V., Shkumatov, A.V. and Chinnathambi, S. (2017) Conformational Dynamics of Intracellular Tau Protein Revealed by CD and SAXS. In Tau Protein ed.^eds), pp. 3–20. Springer

